# Spatiotemporal Mapping of Phosphorylation and Oxidation in the Pig Lens

**DOI:** 10.64898/2026.06.15.732367

**Authors:** Julianna S. Moock, Owen Kelley, Michael Riffle, Gennifer E. Merrihew, Michael J. MacCoss, Jeremy A. Whitson

## Abstract

**Background:** The developmental pattern of the crystalline lens provides a unique model to study biological aging and its effects on the posttranslational modification of long-lived proteins. The orderly differentiation of lens fiber cells leads to a spatiotemporal gradient where mature, organelle-free fiber cells are packed in the lens nucleus surrounded by concentric rings of successively younger fiber cells in the cortex.

**Methods:** Pig lenses were separated into six layers by dissolution in a hypotonic buffer. The changes in protein abundance, oxidation, and phosphorylation that occur across the spatiotemporal gradient of the lens were assessed quantitatively by using data independent acquisition label-free proteomic analysis of these six fractions.

**Results:** Expected changes in protein abundance of major lens protein which reflect the maturation process of lens fiber cells across the spatiotemporal gradient were found. Significant differences were noted in phosphorylation sites on crystallins, phakinin, and actin. Significant changes in oxidation of residues on essential lens proteins, as well as several glycolytic enzymes, were found across the spatiotemporal gradient.

**Conclusion:** Dissolution of the lens followed by high resolution data-independent acquisition proteomics is a powerful technique for spatial mapping of protein abundance and posttranslational modification changes in the lens. The oxidation and phosphorylation sites noted in this study may play important roles in both lens development and cataractogenesis.

## 1. Introduction

Among vertebrate species, the crystalline lens is a widely conserved, transparent tissue of the eye that focuses transmitted light onto the retina with high refractive power using a highly ordered structure of abundant crystallin proteins [1]. The monolayer of cuboidal epithelia of the lens’ anterior surface are the only actively mitotic cells of the lens. As the lens matures, peripheral epithelial cells differentiate into fiber cells that elongate and migrate towards the center of the lens, simultaneously degrading their nucleus and other organelles via proteolytic and apoptotic pathways [2]. The orderly differentiation of fiber cells continues throughout life, leading to development of a spatiotemporal gradient with the oldest denucleated mature fiber cells packed in the lens nucleus and increasingly younger fiber cells in concentric rings of the lens cortex. Given the low abundance of organelles, proteins alone make up 33% of the lens’ wet weight and α and β/γ crystallins comprise nearly 90% of these proteins [3]. While the lack of organelles confers transparency to the lens, it also prohibits cell and protein turnover [4]. Once synthesized, proteins in mature lens fiber cells are never replaced. Thus, the lens nucleus features the same proteins that were present during embryonic development. Alterations to the posttranslational modification (PTM) of these extremely long-lived proteins, including accumulation of damaging nonenzymatic modifications such as oxidation of methionine, are abundant and are believed to be a primary driver of age-related nuclear cataract formation [3]. These PTMs result in alternate proteoforms which may differ in behavior and solubility from the unmodified proteoform, resulting in dysfunction and aggregation. Therefore, analysis of PTMs across the spatiotemporal gradient of the lens is essential to understanding the development of cataracts.

Prior research on the crystalline lens has extracted and lysed whole lenses at various stages of embryonic development to study longitudinal change [5], but this methodology cannot capture long-term accumulation of PTMs. Stewart et al. demonstrated that a single adult lens can be divided into several distinct sections by freezing the lens and shaving concentric layers away from the surface, though they reported slices of the lens nucleus contained fragments of younger cells [6]. Matrix-assisted laser desorption/ionization (MALDI) imaging mass spectrometry (IMS) of frozen sectioned lenses has been demonstrated to be a successful methodology for spatial mapping of lens proteins, although it appears to have limited dynamic range, making it best for analyzing a small set of proteins in a single experiment [7]. As an alternative to these methods, research has shown that gentle stirring in a dissolution buffer is sufficient to separate the cortex from the nucleus of a human lens, given that protein content in solution is associated with the percentage of lens radii removed in solution [8]. Rather than cortex and nucleus alone, the dissolution method can be adapted to divide lens tissue into several sections of equal protein content to observe stepwise changes that have occurred over time. This less intensive separation technique paired with data-independent acquisition (DIA) proteomics provides a more precise method to study age-related PTMs on lens proteins compared to mechanical separation while providing robust data with a high dynamic range.

This study uses porcine (*Sus scrofa domesticus*) lenses due to their widespread availability for study and comparable morphology to human lenses [9]. With this single porcine endpoint tissue, temporal changes in lens fiber cell proteome composition can illuminate specific changes that have occurred due to biological aging. In the present study, we provide proteomics data obtained from six different fractions along the spatiotemporal gradient of the pig lens using DIA to quantify protein abundance as well as two of the most abundant PTMs: oxidation of methionine and phosphorylation of tyrosine, serine, and threonine residues.

## 2. Materials & Methods

### 2.1. Sample Collection

Pig eyes were obtained from a local meat packing site with an approved permit to Obtain Specimens from Official Establishments from the North Carolina Department of Agriculture and Consumer Services Meat and Poultry Inspection Division. Eyes were frozen at -80°C until further use.

### 2.2. Lens Fractionation

Frozen pig eyes were thawed at 4°C. Lenses were removed by carefully cutting into the back of the eye with a scalpel and placed in a sterile Petri dish containing phosphate-buffered saline (PBS). The lens capsule was removed and the remainder of the lens was placed in a well with 6 ml of dissolution media (5 mM Tris, pH 8.0, 1 mM of EDTA, 5 mM of β-mercaptoethanol, 0.2% w/v sodium dodecyl sulfate (SDS)) and a stir bar on a magnetic stir plate set to approximately 350 rpm.

As Mitchell and Cenedella demonstrated in human lenses, dissolution in a hypotonic buffer over a given time will lead to the collection of a consistent percentage of the total lens volume [8]. Here, this method was adapted to porcine lenses across six time intervals to collect multiple sections equal in protein content. As lens radius, and therefore volume, vary between human and porcine lenses, the time for complete dissolution of the lens is also different. To ascertain useful intervals, lens fibers were collected at initial time points of 1, 2, 4, 8, 15, 30, 60, 90, and 120 minutes. Protein content was assayed, and the cumulative protein collected over time was determined for each lens. Using the linear regression of protein dissolved over time, new time points that predicted equal protein dissolution were selected. Protein content was assayed in successive lenses and the subsequent experimental intervals were determined using the linear regression of the combined lenses. As this protocol was repeated, the coefficient of variation of protein obtained at each interval decreased. These methods were replicated until the coefficient of variation approached a constant value at the final chosen time points: 25, 48, 71, 94, 117, and 140 min (Figure A1). Aliquots were analyzed for protein content using a Nanodrop 2000 spectrophotometer (Thermo Fisher Scientific, Waltham, MA). Samples from different pigs served as biological replicates, and timepoints from the same pig were treated as repeated measures. No technical replicate sample preparations were performed.

### 2.3. Sample Preparation and System Suitability

The sample buffer was adjusted to increase SDS to 5% and to add protease and phosphatase inhibitors to each 50 μg of sample. Samples were lysed in a Barocycler 2320 EXT (Pressure Biosciences Inc, Easton, MA) for 30 repeat cycles of 20 sec at 45k psi followed by 10 sec at ambient pressure at 35°C. Lysates were reduced with 20 mM dithiothreitol (DTT), alkylated with 40 mM iodoacetamide (IAA) and quenched with 20 mM DTT. A process control of 800 ng of yeast enolase protein (Sigma-Aldrich, St. Louis, MO) was added to the lysates prior to reduction to monitor digestion. Lysates were prepared for S-trap column (Protifi, Fairport, NY) binding by the addition of 1.2% phosphoric acid and 350 μl of binding buffer (90% methanol, 100 mM tetraethylammonium bromide (TEAB)). The acidified lysate was bound to the column incrementally, followed by 3 wash steps with binding buffer to remove SDS and 3 wash steps with 50:50 methanol:chloroform to remove lipids and a final wash step with binding buffer. Trypsin (1:10) in 50 mM TEAB was added to the S-trap column for digestion at 47°C for one hour. Hydrophilic peptides were eluted with 50 mM TEAB followed by elutions of more hydrophobic peptides with first 0.1% trifluoroacetic acid (TFA) and then 50% acetonitrile. Elutions were pooled, speed vacuumed and resuspended in 0.1% TFA.

An aliquot of ∼15 μg of each digested pool was removed prior to fractionation with a Pierce high pH reversed-phase peptide spin column (Pierce Biotechnology, Waltham, MA; catalog# 84868) to represent the unfractionated sample. The remaining digested pool was loaded into an equilibrated column. The flow-through and water wash was collected. Each fraction collected represents a step gradient of increasing acetonitrile concentrations in a high pH solution. A total of eight fractions is collected for each digested pool. Fractions 1-8, flow- through and the wash were speed vacuumed and resuspended in 0.1% TFA. One μg of each fraction and 150 femtomole of Pierce Retention Time Calibrant (PRTC) were loaded onto the microcapillary trap and C18 column attached to a Thermo EASY nano-flow UHPLC (Thermo Fisher Scientific, Waltham, MA). The PRTC was used to assess quality of the column before and during analysis. We analyze four of these system suitability runs prior to any sample analysis and then after every six to eight sample runs another system suitability run is analyzed. Buffer A is 0.1% formic acid in water and buffer B is 0.1% formic acid in 80% acetonitrile. The 40-minute system suitability gradient consists of a 0 to 16% B in 5 minutes, 16 to 35% B in 20 minutes, 35 to 75% B in 1 minute, 75 to 100% B in 5 minutes, followed by a wash of 9 minutes and a 30-minute column equilibration. The 85-minute sample LC gradient consists of a 0 to 40% B for 60 minutes, 40 to 75% B in 10 minutes, 75 to 100% B in 10 minutes, followed by a 5-minute wash and a 15-minute column equilibration. Peptides are eluted from the column with a 50°C heated source (CorSolutions, Ithica, NY) and electrosprayed into a Thermo Orbitrap Fusion Lumos Mass Spectrometer with the application of a distal 3 kV spray voltage. For the system suitability analysis, a cycle of one 120,000 resolution full-scan mass spectrum (350-2000 m/z) followed by a data-independent MS/MS spectra on the loop count of 76 data-independent MS/MS spectra using an inclusion list at 15,000 resolution, AGC target of 4e5, 20 millisecond (ms) maximum injection time, 33% normalized collision energy with a 8 m/z isolation window. Each DDA fraction run consists of a cycle of one 60,000 resolution full-scan orbitrap mass spectrum with a mass range of 400-1600 m/z, AGC target of 4e5, 50 ms maximum injection time followed by a data-dependent MS/MS spectra with “rapid” ion trap scanning, AGC target of 1e4, quadrupole isolation width of 1.6 m/z and 35 ms maximum injection time. The MS/MS spectra were acquired using 27% normalized collision energy (NCE), allowing precursor charge states of 2-7 and dynamic exclusion was set for 20 s using monoisotopic precursor selection (MIPS) with a mass error of 10 ppm. Application of the mass spectrometer and LC solvent gradients are controlled by the ThermoFisher Xcalibur data system.

System suitability runs were analyzed using Skyline [10,11] and AutoQC [12] via PanoramaWeb [13]. Thermo XCalibur RAW files were converted to mzML format using Proteowizard [14] using vendor peak picking and searched using Comet [15] with a 15 ppm precursor tolerance, fixed modification of 57.02 C and Percolator [16] with a q-value of 0.01 against a FASTA database containing protein sequences from domestic pig (reviewed and unreviewed combined – UniProt [17] ID UP000008227) and contaminants to generate a sub-database.

### 2.4. DIA Analysis of Lens Protein Samples

The 110-minute DIA sample LC gradient consists of a 2 to 7% B for 1 minutes, 7 to 14% B in 35 minutes, 14 to 40% B in 55 minutes, 40 to 60% B in 5 minutes, 60 to 98% B in 5 minutes, followed by a 9-minute wash and a 30-minute column equilibration. Peptides were eluted from the column with a 50°C heated source (CorSolutions, Ithica, NY) and electrosprayed into a Thermo Orbitrap Fusion Lumos Mass Spectrometer with the application of a distal 3 kV spray voltage. For the system suitability analysis, a cycle of one 120,000 resolution full-scan mass spectrum (350-2000 m/z) followed by a data-independent MS/MS spectra on the loop count of 76 data-independent MS/MS spectra using an inclusion list at 15,000 resolution, AGC target of 4e5, 20 millisecond (ms) maximum injection time, 33% normalized collision energy with an 8 m/z isolation window. For the sample digest, first a chromatogram library of six independent injections were analyzed from a pool of all samples within the first batch. For each injection a cycle of one 120,000 resolution full-scan mass spectrum with a mass range of 100 m/z (400-500 m/z, 500-600 m/z, 600-700 m/z, 700-800 m/z, 800-900 m/z, 900-1000 m/z) followed by a data-independent MS/MS spectra on the loop count of 26 at 30,000 resolution, AGC target of 4e5, 60 ms maximum injection time, 33% normalized collision energy with a 4 m/z overlapping isolation window. The chromatogram library data was used to quantify proteins from individual sample runs. These individual runs consist of a cycle of one 120,000 resolution full-scan mass spectrum with a mass range of 350-2000 m/z, AGC target of 4e5, 100 ms maximum injection time followed by a data-independent MS/MS spectra on the loop count of 76 at 15,000 resolution, AGC target of 4e5, 20 ms maximum injection time, 33% normalized collision energy with an overlapping 8 m/z isolation window. Application of the mass spectrometer and LC solvent gradients are controlled by the Thermo Fisher Xcalibur data system. No technical replicate LC–MS/MS injections were performed.

System suitability and process control runs were analyzed using Skyline [10,11] and AutoQC [12] via PanoramaWeb [13]. Thermo XCalibur RAW files were converted to mzML format using Proteowizard [14] using vendor peak picking and demultiplexing with the settings of “overlap_only” and Mass Error = 10.0 ppm. On-column chromatogram libraries were created using the data from the six gas phase fractionated “narrow window” DIA runs of the pooled samples from each batch. These narrow windows were analyzed using EncyclopeDIA [18–21] with the default settings (10 ppm tolerances, trypsin digestion, HCD b- and y-ions) of a Prosit [22] predicted spectra library based the UniProt pig (reviewed and unreviewed combined) FASTA. The results from this analysis were saved as a “Chromatogram Library”; in EncyclopeDIA’s eLib format where the predicted intensities and iRT of the Prosit library were replaced with the empirically measured intensities and RT from the gas phase fractionated LC-MS/MS data. The “wide window” DIA runs were analyzed using EncyclopeDIA requiring a minimum of 3 quantitative ions and filtering peptides with q-value ≤ 0.01 using Percolator 3.01. After analyzing each file individually, EncyclopeDIA was used to generate a “Quant Report”; which stores all the detected peptides, integration boundaries, quantitative transitions, and statistical metrics from all runs in an eLib format. The Quant Report eLib library was imported into Skyline with the pig UniProt FASTA as the background proteome to map peptides to proteins, perform peak integration, and normalize to total ion current (TIC). A csv file of modified and unmodified peptide level total area fragments (TAFs) for each replicate was exported from Skyline using the custom reporting capabilities of the document grid.

### 2.5 Generation of a Carafe Fine-Tuned Spectral Library

A single data-independent acquisition (DIA) run was used to fine-tune the Carafe fragment intensity and retention time models (11May2022-Lumos-DIA-Whitson-Porcine-Lens-B1-ind-8mz-ovlp-400to1000-PL07), a porcine lens tryptic digest acquired on a Thermo Orbitrap Fusion Lumos using overlapping 8 m/z isolation windows across a precursor range of 400 to 1000 m/z. This run was selected as a single representative file of moderate size from the middle of the study. Spectra were processed with msconvert (ProteoWizard) using the demultiplexing [23] and SIM-as-spectra options to resolve the overlapping isolation windows.

The processed DIA data were searched with DIA-NN (version 1.9.2) in library-free mode against the UniProt *Sus scrofa* canonical proteome (downloaded November 2024), supplemented with common contaminants and yeast enolase 1 (ENO1). Trypsin specificity was applied (cleavage rule K*,R*,!*P) allowing up to one missed cleavage. Peptides of 7 to 30 residues were considered, with precursor m/z from 400 to 1000, precursor charge 2 to 3, and fragment m/z from 200 to 2000. Carbamidomethylation of cysteine was set as a fixed modification. Oxidation of methionine (UniMod:35) and phosphorylation of serine, threonine, and tyrosine (UniMod:21) were allowed as variable modifications, with at most two variable modifications per peptide. Retention time profiling was enabled, protein grouping was set to pg-level 1, and identifications were filtered to a 1% precursor q-value.

The DIA-NN search results were then supplied to Carafe (version [1.0.0]) to fine-tune its fragment intensity and retention time predictors and to generate an in silico spectral library against the same FASTA. Carafe was run in phosphorylation mode. Carbamidomethylation of cysteine was specified as a fixed modification. Oxidation of methionine together with phosphorylation of serine, threonine, and tyrosine were declared as variable modifications in a single combined argument (varMod 2,7,8,9), allowing up to three variable modifications per peptide. The predicted fragment m/z range was 200 to 2000.

Consistent with the upstream search, the predicted library was restricted to precursor charges 2 to 3, since broader charge ranges (1 to 4) exhausted available memory during prediction.

Spectrum conversion, the DIA-NN search, and Carafe fine-tuning and library prediction were carried out with the nf-carafe-ai-ms Nextflow pipeline [24]. Jobs were run locally with a ceiling of 30 GB of memory and 14 CPU cores.

### 2.6 Gas-Phase Fractionated Library Refinement

The Carafe predicted library was empirically refined against gas-phase fractionated (GPF) data to restrict it to peptides detectable in the sample matrix, without altering the predicted fragment intensities or retention times. Six narrow-window GPF runs collectively spanning 400 to 1000 m/z (400 to 500, 500 to 600, 600 to 700, 700 to 800, 800 to 900, and 900 to 1000 m/z), each acquired with overlapping 4 m/z isolation windows on the Orbitrap Fusion Lumos, were searched with DIA-NN (version 1.9.2) using the Carafe library as input. Per-run mass accuracy and isolation window settings were determined individually (--individual-mass-acc, --individual-windows). Trypsin specificity was ap-plied (K*,R*) with up to one missed cleavage, carbamidomethylation of cysteine as a fixed modification, and oxidation of methionine and phosphorylation of serine, threonine, and tyrosine as variable modifications (up to two per peptide). Conservative machine learning mode and peptidoform scoring were enabled, protein inference was relaxed (--relaxed-prot-inf, pg-level 1), normalization was global, and results were filtered to a 1% q-value. A new spectral library was generated from the search (--gen-spec-lib) and reannotated against the same *Sus scrofa* FASTA. DIA-NN optimized mass accuracy to 7 ppm at both MS1 and MS2. The refined library contained 1,721 protein groups (1,901 protein isoforms) and 13,817 precursors (13,676 target precursors).

### 2.7 DIA Quantification of Individual Samples

The GPF-refined library was used to quantify all individual-sample DIA runs (samples PL01 through PL36, plus pooled and replicate injections, acquired across three batches) collected with staggered 8 m/z isolation windows spanning 400 to 1000 m/z. DIA-NN (version 1.9.2) was run with peptides of 7 to 30 residues, precursor m/z 400 to 1000, precursor charge 2 to 3, trypsin specificity (K*,R*) with one missed cleavage, fixed carbamidomethylation of cysteine, and variable oxidation of methionine and phosphorylation of serine, threonine, and tyrosine (up to two per peptide). Conservative machine learning mode and peptidoform scoring were enabled, and protein inference used heuristic grouping (--relaxed-prot-inf). The analysis was run in double-pass mode (--reanalyse), in which DIA-NN builds a spectral library from the runs and uses it to reprocess them, retaining the input library spectra with empirically aligned retention times. Mass accuracy was optimized automatically from the first run, retention time profiling and global normalization were applied, and precursors and protein groups were filtered to a 1% q-value with quantitative matrices exported. The resulting spectral library (report-lib.parquet.skyline.speclib) was imported into Skyline [25,26] for downstream quantification, peak boundary imputation [27], and report generation.

### 2.8. Identification of Time-Associated Proteins by Generalized Estimating Equations

Raw protein abundance output from Skyline was loaded using pandas [28] and NumPy [29], with missing values encoded as zeros and a pooled quality-control sample excluded from statistical modeling. Proteins with more than 50% zero values across samples were removed; the remaining samples were normalized by L1 (total-signal) scaling with the ‘L1Normalizer‘ from the ‘pronoms‘ Python package (https://github.com/mriffle/pronoms) so that each sample’s total protein signal was equal, and systematic inter-batch variation was then removed with the empirical-Bayes ComBat algorithm [30] implemented in ‘pycombat‘ (https://github.com/epigenelabs/pyComBat) [31], using the ‘Batch‘ metadata column as the batch variable. The batch-corrected matrix was offset log-transformed using ‘numpy.log1p‘, and for each protein a Generalized Estimating Equation (GEE) [32] model of the form ‘protein_value ∼ timè was fit with ‘stats-models.formula.api.geè (https://www.statsmodels.org) [33] using a Gaussian family, identity link, and an exchangeable working correlation structure with ‘Pig‘ as the cluster-ing variable (so that all within-pig sample pairs share a common correlation), where ‘protein_valuè is the per-protein abundance standardized to zero mean and unit variance across samples and ‘timè is the timepoint treated as a continuous predictor; samples were sorted by pig and timepoint prior to fitting, proteins with fewer than three complete observations or fewer than two distinct pigs were skipped, and the coefficient, robust (sandwich) standard error, Wald p-value, and 99% Wald confidence interval for the ‘timè term were retained for each fitted model, with raw p-values adjusted across proteins by the Benjamini–Hochberg procedure [34] via ‘statsmodels.stats.multitest.multipletests‘ and proteins with BH-adjusted q < 0.01 declared significant; p-values were computed under the normal approximation using ‘scipy.stats‘ [35]. For visualization only, Wald p-values of exactly zero (floating-point underflow) were replaced with half of the smallest non-zero observed p-value so that −log₁₀(p) remained finite, and results were displayed as a volcano plot of the GEE coefficient for ‘timè against −log₁₀(q) drawn with ‘matplotlib‘ (https://matplotlib.org) [36], with a reference line at q = 0.01 and the 15 most significant proteins labeled using ‘adjustText‘ (https://github.com/Phlya/adjustText) to resolve label collisions.

### 2.9. Principal Component Analysis of Peptide Abundance vs. Timepoint

Raw peptide abundance output from Skyline was loaded using pandas and NumPy, with missing values encoded as zeros. Duplicate peptide entries (same modified peptide sequence mapped to multiple protein groups) were collapsed to a single column, with the associated protein labels merged as a comma-delimited string. Peptides with more than 50% zero values across samples were removed; remaining samples were normalized by per-sample median scaling using the ‘MedianNormalizer‘ from the ‘pronoms‘ Python package so that each sample’s median peptide intensity was equal across samples, and systematic inter-batch variation was removed with ‘pycombat‘, as described above. Principal component analysis (PCA) was performed on the normalized, batch-corrected peptide matrix after excluding the pooled quality-control samples; features were centered and scaled to unit variance with ‘sklearn.preprocessing.StandardScaler‘ and the first four principal components were computed with ‘sklearn.decomposition.PCÀ (https://scikit-learn.org) [37]. Samples were projected into two scatter plots (PC1 vs. PC2 and PC3 vs. PC4) drawn with ‘matplotlib‘. For each PCA panel, the axes of each scatter plot are flanked by marginal plots that summarize the relationship between that principal component and timepoint: the top-marginal panel plots the PC1 (or PC3) score on the x-axis against timepoint on the y-axis, and the right-marginal panel plots the PC2 (or PC4) score on the y-axis against timepoint on the x-axis, with points colored by timepoint using the same colormap as the main scatter. An ordinary least-squares (OLS) regression line was fit between each PC score and timepoint using ‘scipy.stats.linregress‘ (https://scipy.org) [35]; the fitted line is overlaid on the marginal scatter with a shaded band indicating the 95% confidence interval of the mean response, and the panel is annotated with the regression slope and its two-sided p-value (the Wald test that the slope differs from zero). Spearman rank correlations between each PC score and timepoint were computed in parallel with ‘scipy.stats.spearmanr‘ as a non-parametric check of monotonic association.

### 2.10. Functional Categorization of Lens Proteins and Per-Category Heatmaps

Protein functional categories of interest were assigned from the same FASTA file used for proteomics analysis by parsing each entry’s header to extract the accession, description, organism, taxonomy identifier (OX), and gene name (GN). Only entries with OX = 9823 (*Sus scrofa*) were retained. Each retained protein was assigned to zero or more curated functional categories by matching case-insensitive keyword and regular-expression rules against the protein description, the UniProt gene name, and gene-like tokens extracted from the description. Protein quantitation was loaded, missingness-filtered, L1-normalized, and ComBat batch-corrected exactly as described in the GEE section above. For each category, proteins present in both the category mapping and the batch-corrected matrix were retained, abundances were averaged across replicates within timepoint using ‘pandas.DataFrame.groupby‘ [28], and each protein row was standardized across timepoints by a z-score (‘scipy.stats.zscorè with ‘nan_policy=’omit’‘) [35]. Heatmaps were drawn with ‘seaborn.heatmap‘ (https://seaborn.pydata.org) [38].

### 2.11. Peptide-Level GEE and Identification of Modification-Discordant Peptide Pairs

A peptide-level analogue of the protein GEE analysis described above was run. Peptides sharing an identical modified sequence across multiple protein groups were collapsed into a single column, peptides were then missingness-filtered at the 50% threshold, and L1 (total-signal) normalized with the ‘pronoms‘ ‘L1Normalizer‘. The normalized ma-trix was log1p-transformed (‘numpy.log1p‘), pooled QC samples were excluded, and a GEE model of the form ‘peptide_value ∼ timè was fit per peptide with ‘statsmodels.formula.api.geè using a Gaussian family, identity link, and an exchangeable working correlation structure clustered on ‘Pig‘, exactly as described in the protein GEE section. For each post-translational modification of interest, specified as an integer delta-mass (+16 Da, consistent with oxidation; +80 Da, consistent with phosphorylation), peptides were paired by base sequence and the respective modified form. A pair was retained as "discordant" when the modified form was statistically significant (BH-FDR ≤ 0.05 on the ‘timè term, non-zero coefficient) and the unmodified form either (i) failed the significance threshold or (ii) passed the significance threshold but had a coefficient of the opposite sign. When multiple candidate pairs existed for a base peptide, a single representative pair was se-lected by maximizing |coef_mod − coef_unmod| (absolute discordance), breaking ties on the smaller FDR of the modified form.

### 2.12. Traditional Differential Abundance Analysis of Proteins and Peptides

Raw protein and peptide abundance output from Skyline was loaded using pandas and NumPy, with missing values encoded as zeros. Samples were normalized by L1 (to-tal-signal) scaling with the ‘L1Normalizer‘ from the ‘pronoms‘ Python package (https://pypi.org/project/pronoms/), and the per-sample/per-protein missingness mask (zero or NaN on the L1-normalized matrix) was recorded for later restoration. Systematic inter-batch variation was removed with Combat (as previously described) and the recorded missingness mask was re-applied as ‘NaN‘ so that ComBat-induced shifts at originally-missing cells did not contribute to downstream fitting. The pooled quality-control sample was excluded from all statistical modeling. For each timepoint comparison, the batch-corrected matrix was restricted to the two timepoints of interest and offset log2-transformed (‘numpy.log2(1 + x)‘) to put abundances on the scale expected by limma. Pigs were used as the pairing/blocking unit and a binary class indicator (0 = earlier timepoint, 1 = later timepoint) was assigned per sample. Differential abundance was tested with the limma empirical-Bayes moderated linear model [39,40] from the Bioconductor ‘limmà package, called from Python via ‘rpy2‘ (https://rpy2.github.io /) with R 4.4.2 [41].Within R, the design matrix ‘∼ subject + class‘ was constructed with ‘model.matrix‘, where ‘subject‘ is a factor of pig identifiers and ‘class‘ is the numeric timepoint indicator; protein-wise linear models were fit with ‘limma::lmFit‘ and moderated with ‘limma::eBayes‘, and per-protein log2 fold-changes, p-values, and Benjamini–Hochberg-adjusted q-values [34] for the ‘class‘ coefficient were extracted. NaN values in the input matrix were passed through to ‘lmFit‘, which fits each protein on its complete observations.

## 3. Results

### 3.1. Protein Abundance Changes Across the Lens Spatiotemporal Gradient

The results were analyzed longitudinally using the Generalized Estimating Equation (GEE) to determine how the abundance of each protein and peptide (modified and un-modified) changes across the spatiotemportal gradient. Figure 1A compares the GEE regression coefficient vs. -log of the Benjamini-Hochberg FDR-corrected P-values for protein abundances. Of the 754 proteins which were analyzed, 52 showed significantly increasing abundance and 350 showed significantly decreasing abundance moving from the lens periphery to the nucleus. Principle component analysis (PCA) showed clear clustering of the data based upon timepoint of dissolution and, therefore, location in the spatiotemporal gradient of the lens (Figure 1B). Pantherdb.org was used to analyze the impact of post-translational modifications on biological processes (PANTHER GO-Slim Biological Process) in the dataset by performing a statistical overrepresentation test for the list of proteins which showed significant (P<0.05) increase or decrease across the spatiotemportal gradient based on the GEE (Figure 1C). Unsurprisingly, given the unique and highly specialized protein milieu of the lens, processes such as lens development in camera-type eye, eye development, and visual perception were significantly overrepresented. This is mostly owed to the high abundance of crystallins and other unique lens proteins, such as beaded filament structural proteins, in this proteomic dataset. Similarly, actin is a major component of the lens cytoskeleton and essential to morphological changes that occur as part of the fiber cell differentiation process [42] and mature lens fiber cells must rely entirely on glycolysis for energy metabolism [43], so actin filament depolymerization and glycolytic process being major affected pathways across the spatiotemporal gradient are expected results.

**Figure 1.**
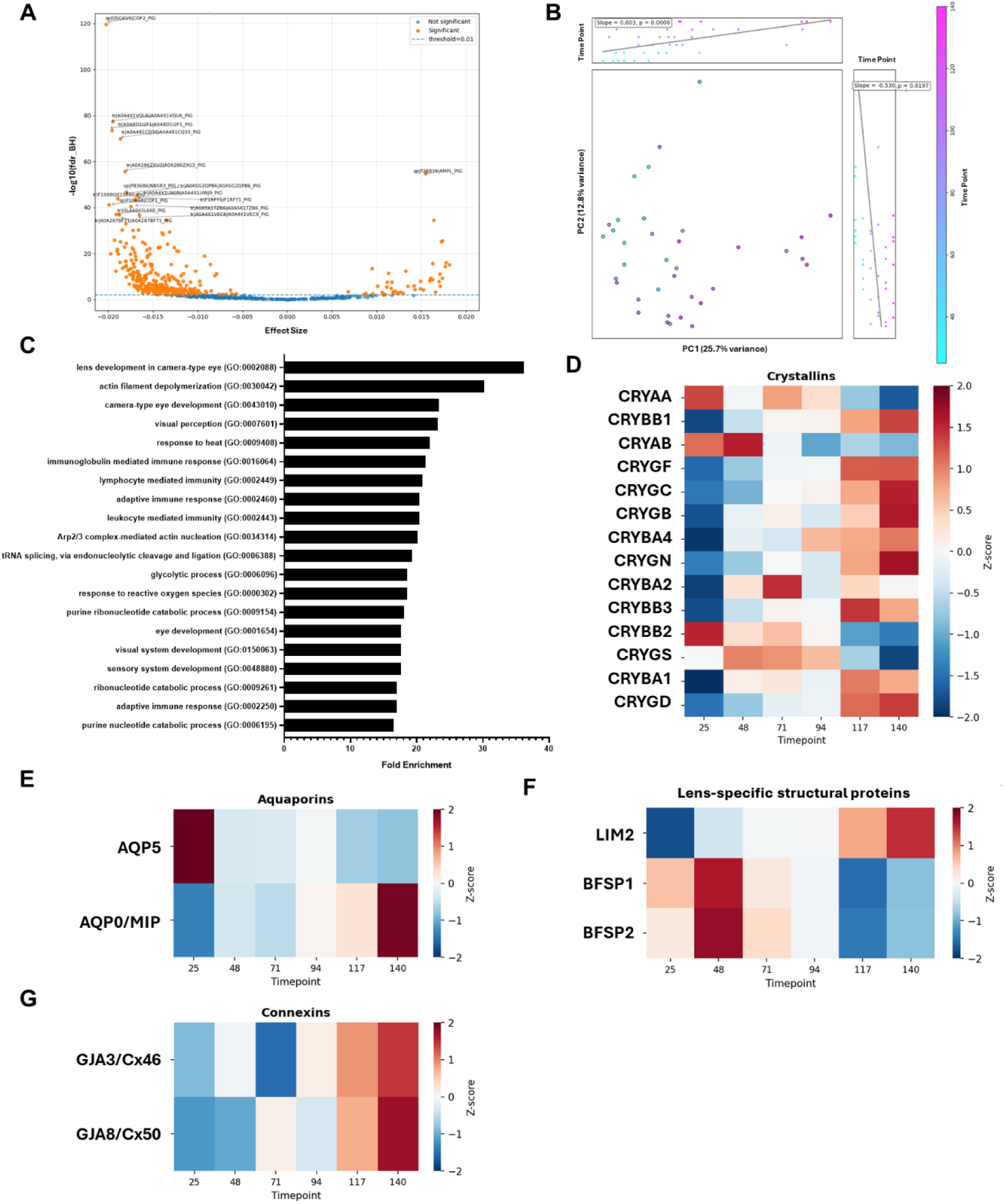
Overview of Trends in Protein Abundance Across the Lens Spatiotemporal Gradient. (**A**) GEE plot of regression coefficient vs. -log of the Benjamini-Hochberg FDR-corrected P-values. Each point represents a unique protein. Blue points do not reach the threshold of significance while orange points are statistically significant. **B**) Principal component analysis plot. (**C**) Top 20 statistically overrepresented biological processes (GO-slim terms) in the list of significant (P<0.05) proteins via pantherdb analysis. Results of the analysis are ordered by fold-enrichment. **(D-G)** Heatmaps showing the relative abundance of major lens proteins across the spatiotemporal gradient. All timepoints n=6.

Heatmaps showing the relative abundance of essential lens proteins in categories of crystallins (Figure 1D), aquaporins (Figure 1E), lens-specific structural proteins (Figure 1F), and connexins (Figure 1G) are shown. With the exception of αA crystallin (CRYAA), αB crystallin (CRYAB), and βB2 crystallin (CRYBB2), crystallin abundances increased across the spatiotemporal gradient when moving from the periphery toward the nucleus (Figure 1D). Aquaporin 5 (AQP5) and aquaporin 0 (AQP0) levels showed an inverse relationship, with AQP5 decreasing in abundance going from the periphery to the nucleus and AQP0 increasing in abundance moving along the same axis (Figure 1E), matching the pattern which has been well-established in prior studies [44]. Lens intrinsic membrane protein 2 (LIM2) increased in abundance moving towards the nucleus while filensin (BFSP1) and phakinin (BFSP2) decreased (Figure 1F). Connexin 46 (Cx46) and connexin 50 (Cx50) both increased going from the periphery to the nucleus, while connexin 43 (Cx43) was not detected (Figure 1G), which was expected as it is found primarily in lens epithelia and not fiber cells [45]. The full set of protein abundance results, including traditional differential abundance analysis comparing deeper layers to the T32 fraction, can be viewed in Appendix B1.

### 3.2. Analysis of Modified Peptides

In order to determine the site-specific PTMs that are associated with lens fiber cell age, the list of modified peptides which showed significant increase or decrease in abundance across the spatiotemporal gradient of the lens based on the GEE was further filtered to select only those modified peptides whose matching unmodified form did not show a significant change in abundance or showed a significant change in the opposite direction. This eliminates the possibility that any differences detected in the abundance of a modified peptide are simply due to overall differences in the abundance of the peptide rather than a difference in the level of modification. Phosphorylated peptides that demonstrated an increased occupancy with age/location going from the periphery to the nucleus which was discordant with the matching unmodified peptides mapped to sites on actin, BFSP2, αA CRYAA, CRYAB, βA2 crystallin (CRYBA2), and CRYBB2 (Figure 2B-I; Table 1). There were no discordantly altered phosphorylated peptides which decreased in abundance going from the periphery to the nucleus. Statistically overrepresented pathways for the set of proteins with significant changes in phosphorylation were largely similar to what was found based on protein abundance, although there were also a number of pathways related to apoptosis that were noted (Figure 2A).

**Figure 2.**
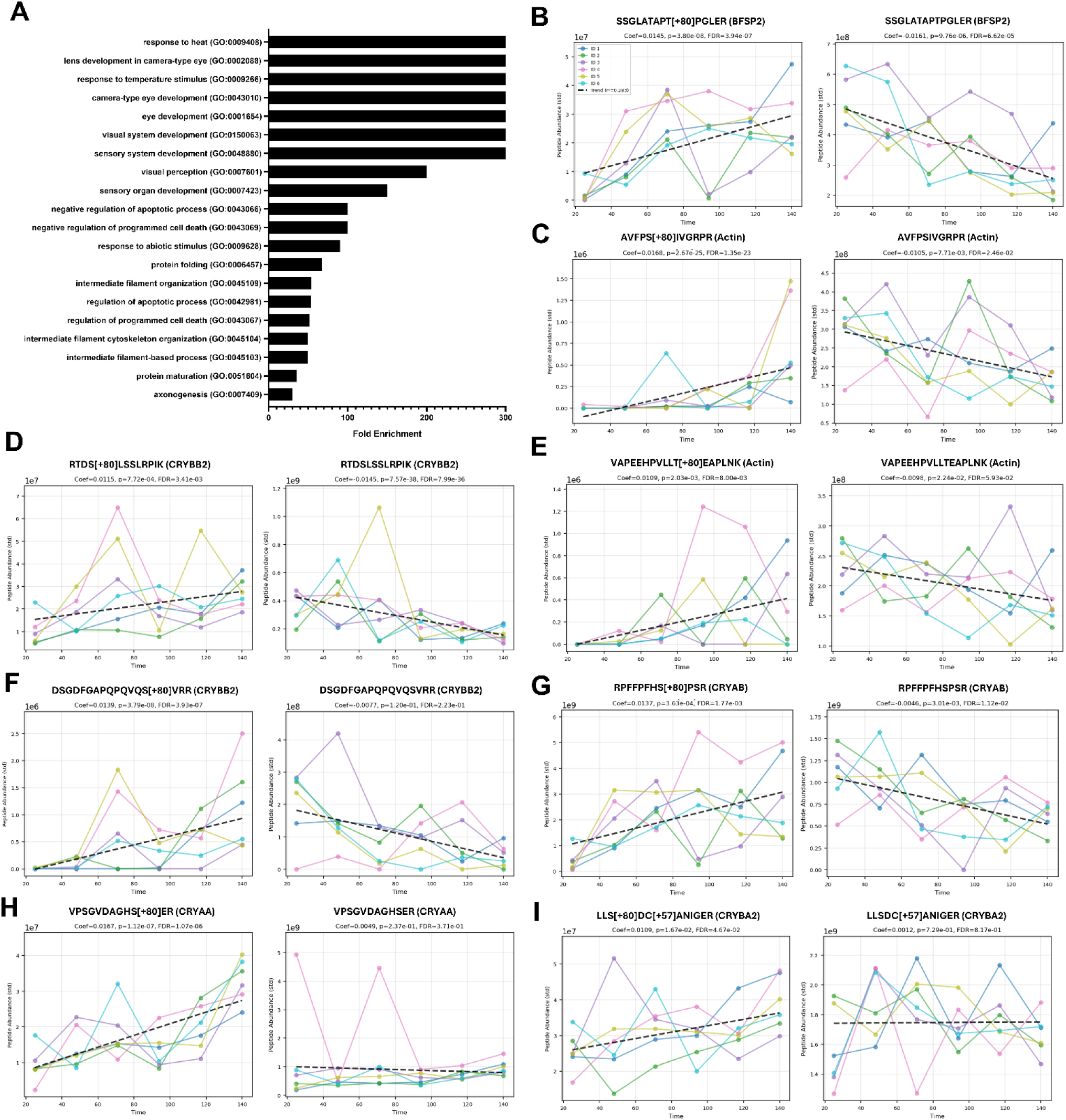
Analysis of Changes in Phosphorylation Across the Spatiotemporal Gradient of the Lens. (**A**) Top 20 statistically overrepresented biological processes (GO-slim terms) in the list of significant (P<0.05) proteins via pantherdb analysis. Results of the analysis are ordered by fold-enrichment. (**B-I**) X-Y plots of peptide abundance for the top 8 discordant phosphorylated peptides and the matching unphosphorylated peptides. Dashed regression lines along with their coefficients, P-values, and FDR-adjusted P-values are shown.

**Table 1.**
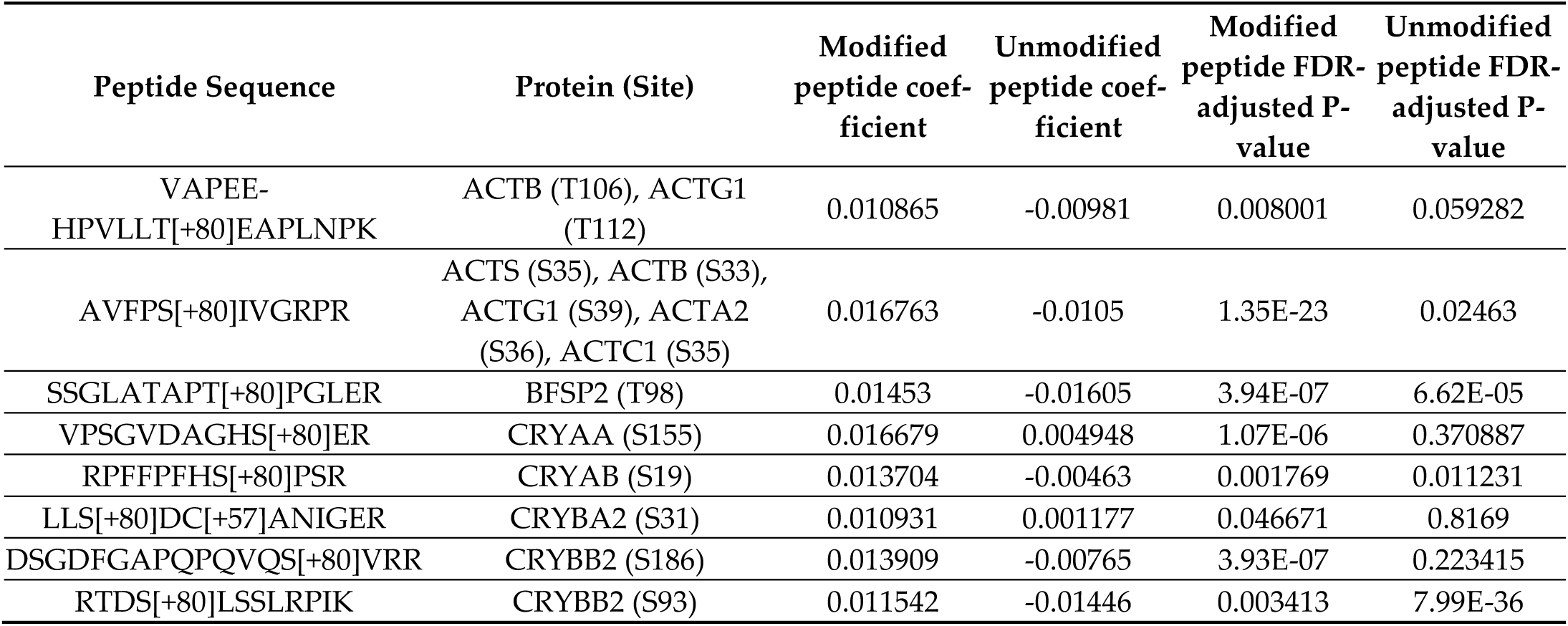
List of Significant Discordantly Altered Phosphorylated Peptides. Coefficients are the slope of the regression line analyzing peptide abundance vs. time. N = 6.

Oxidized peptides that demonstrated a decreased occupancy with age/location moving towards the lens nucleus that was discordant with matching unmodified peptides mapped to sites on quinone oxidoreductase/taxon-specific crystallin Z (QOR/CRYZ), filamin A (FLNA), coactosin-like protein (COTL1), ubiquitin specific peptidase 14 (USP14), karyopherin subunit beta-1 (KPNB1), N-myc downstream-regulated gene 1 (NDGR1), phosphoglycerate mutase 1 (PGAM1) (Figure 3B-H; Table 2), as well as dihydropyrimidinase like 2 (DPYSL2), vimentin (VIM), pyruvate kinase (PKM), phakinin (BFSP2), protein S100-A4 (S100A4), cytoplasmic dynein 1 heavy chain 1 (DYNC1H1), trans-1,2-dihydro-benzene-1,2-diol dehydrogenase (DHDH) (Table 2). Oxidation sites that demonstrated increased occupancy in the direction of periphery to nucleus were found on glucose-6-phosphate isomerase (G6PI) (Figure 2I; Table 2) as well as αB crystallin (CRYAB) and βB1 crystallin (CRYBB1) (Table 2). Statistically overrepresented pathways for the set of proteins with significant changes in oxidation had some overlap with pathways that were overrepresented in the overall protein abundance data, such as visual perception, but were largely dominated by pathways related to energy metabolism, such as ATP metabolic process and carbohydrate metabolic process (Figure 3A).

**Figure 3.**
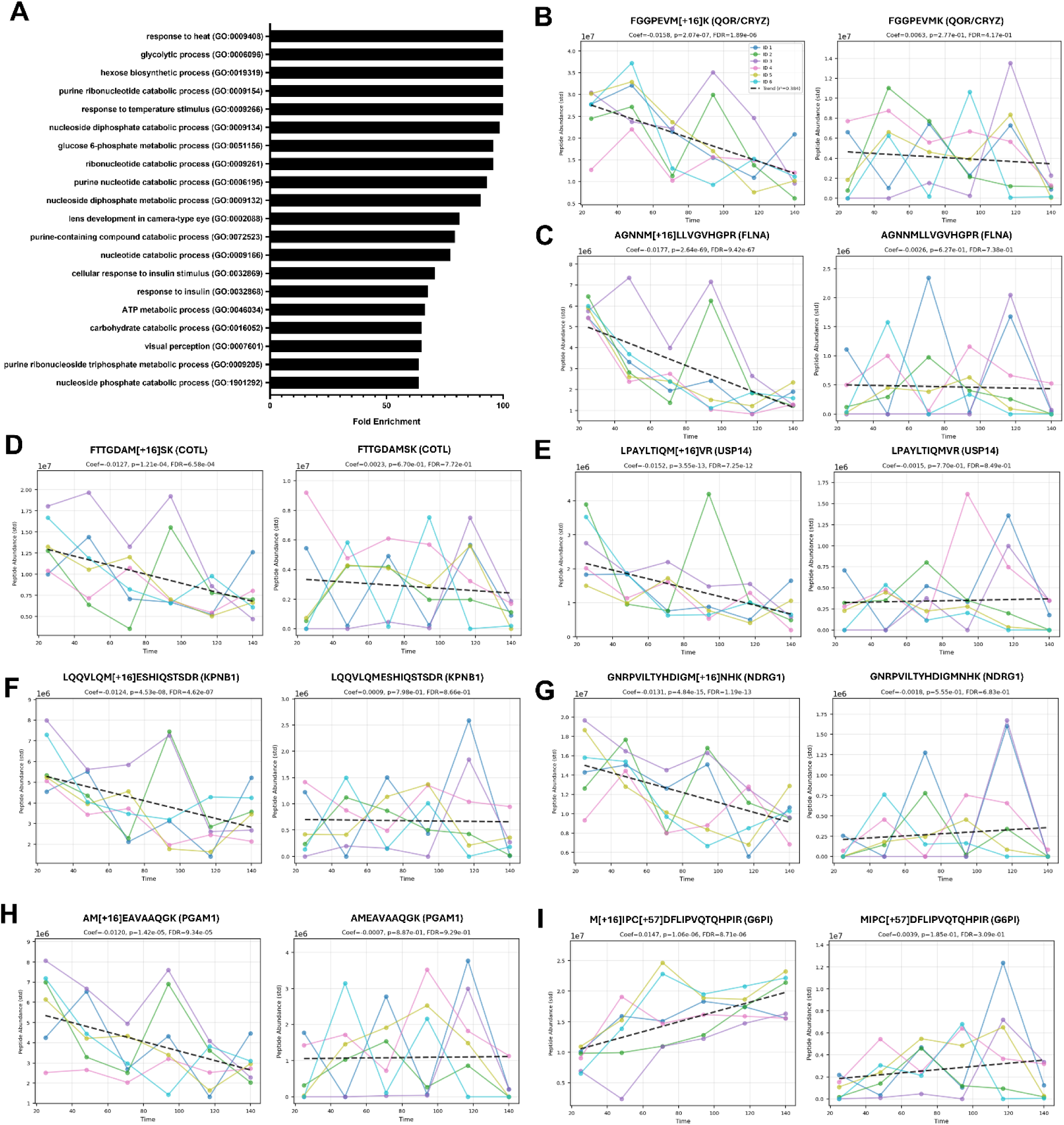
Analysis of Changes in Oxidation Across the Spatiotemporal Gradient of the Lens. (**A**) Top 20 statistically overrepresented biological processes (GO-slim terms) in the list of significant (P<0.05) proteins via pantherdb analysis. Results of the analysis are ordered by fold-enrichment. (**B-I**) X-Y plots of peptide abundance for the top 8 discordant oxidized peptides and the matching un-oxidized peptides. Dashed regression lines along with their coefficients, P-values, and FDR-adjusted P-values are shown.

**Table 2.** List of Significant Discordantly Altered Oxidized Peptides. Coefficients are the slope of the regression line analyzing peptide abundance vs. time. N = 6.

| Peptide Sequence | Protein (Site) | Modified peptide coefficient | Unmodified peptide coefficient | Modified peptide FDR-adjusted P-value | Unmodified peptide FDR-adjusted P-value |
| --- | --- | --- | --- | --- | --- |
| FGGPEVM[+16]K | CRYZ/QOR (M22) | -0.01584 | 0.00626 | 1.89E-06 | 0.416649 |
| AGNNM[+16]LLVGVHGPR | FLNA (M2621) | -0.01766 | -0.00259 | 9.42E-67 | 0.737836 |
| FTTG DAM[+16]SK | COTL1 (M70) | -0.01268 | 0.002278 | 0.000658 | 0.772012 |
| LPAYLTIQM[+16]VR | USP14 (M328) | -0.01521 | -0.00152 | 7.25E-12 | 0.84877 |
| LQQVLQM[+16]ESHIQSTSDR | KPNB1 (M564) | -0.01239 | 0.000909 | 4.62E-07 | 0.866319 |
| GNRPVILTYHDIGM[+16]NHK | NDRG1 (M67) | -0.01313 | -0.00181 | 1.19E-13 | 0.68345 |
| AM[+16]EAVAAQ GK | PGAM1 (M243) | -0.01202 | -0.00073 | 9.34E-05 | 0.929353 |
| M[+16]IPC[+57]DFLIPVQTQHPI<br>R | G6PI (M401) | 0.01471 | 0.003881 | 8.71E-06 | 0.309068 |
| M[+16]VIPGGIDVHTR | DPYSL2 (M169) | -0.0085 | 0.002217 | 0.007347 | 0.766889 |
| LGDLYEEM[+16]R | VIM (M154) | -0.01671 | -0.00703 | 4.35E-41 | 0.336625 |
| RFDEILEASDGIM[+16]VAR | PKM (M291) | -0.00825 | 0.001178 | 0.044161 | 0.86923 |
| RVVVDVPTGASPSM[+16]PLQR | BFSP2 (M18) | -0.01625 | -0.00744 | 4.65E-06 | 0.291146 |
| ALDVM[+16]VSTFHK | S100A4 (M12) | -0.01327 | -0.00478 | 2.65E-05 | 0.486099 |
| LM[+16]LHTENIQAGADDFKER | BFSP2 (M175) | -0.00675 | -0.00044 | 0.047231 | 0.938081 |
| APSWIDTGLSEM[+16]R | CRYAB (M68) | 0.010915 | 0.005621 | 0.011574 | 0.241055 |
| M[+16]VVLSLPR | DYNC1H1 (M989) | -0.00975 | -0.00447 | 2.95E-08 | 0.442239 |
| GLFLM[+16]EAIWTR | DHDH (M121) | -0.01185 | -0.00712 | 0.012536 | 0.321094 |
| HWNDWGAFQPQM[+16]QAVR | CRYBB1 (M223) | 0.009423 | 0.007526 | 0.043383 | 0.196144 |

The full set of peptide abundance results, including traditional differential abundance analysis comparing deeper layers to the T32 fraction, can be found in Appendix B2.

## 4. Discussion

Protein abundance data matched expectations for how different essential lens proteins should change across the spatiotemporal gradient, essentially showing how the proteome of fiber cells changes as they go through the maturation process progressing from the lens periphery to the organelle-free zone. This indicates the success of the dissolution method for the fractionation and proteomic analysis of lenses.

### 4.1. Phosphorylation of Critical Lens Proteins is Altered Across the Spatiotemporal Gradient

Phosphorylation of lens proteins is known to be a critical regulatory mechanism in lens development and maintenance and many phosphosites have been established in the literature. However, significant mystery remains around the exact impacts of phosphorylation on most of these sites.

It is well established that α crystallin function is regulated by phosphorylation, most of which is the result of cAMP-dependent pathways [46,47]. Phosphorylation of the S155 site on CRYAA has previously been observed in bovine lenses [48], as well as phosphorylation of the orthologous S178 site in mouse lenses [49]. The effects of phosphorylation on this site in regard to protein function have not been annotated. Intriguingly, humans do not have an orthologous phosphosite as they have an alanine residue, rather than a serine residue, at the same locus. Phosphorylation of the S19 site on CRYAB is also very well-established but it has been demonstrated that it has little effect on the chaperone function of the protein, whereas phosphorylation of S45, which was not significantly altered in the present analysis, massively alters protein function and lens proteostasis [50]. Although phosphorylation of β crystallins has long been noted in the literature, its effects on their function in the lens remains unclear. However, outside of the lens, phosphorylation of β crystallins has been shown to regulate anti-apoptotic signaling in bovine retinal pigmented epithelia exposed to light-induced oxidative stress [51]. Phosphorylation of CRYBA2 has been found to be extensive in human lenses, with ∼20% of the protein being phosphorylated at the same S31 site noted in this study [52], but the exact consequences of this phosphorylation event have not been determined. CRYBB2 has long been known to contain phosphosites in bovine lenses [53], though the exact loci have not been reported. Phosphorylation at the S186 site of CRYBB2 was previously reported in a study of ischemia in mouse tumors [54], although the role of CRYBB2 in that context and the effect of its phosphorylation remains unclear. To the authors’ knowledge, no publications have reported any data regarding the S93 phosphosite of CRYBB2, which showed significant differences across the spatiotemporal gradient in this study. Overall, the effects of phosphorylation on CRYBB2 remain unknown.

The T98 phosphorylation site found on porcine BFSP2 is orthologous to the S100 phosphorylation site found previously in bovine lenses [55] based upon sequence homology. This conservation across species indicates importance of this phosphosite. However, it should be noted that humans and rodents have a glycine residue, rather than a threonine or serine, at this site making phosphorylation impossible. Therefore, this phosphosite may be specific to order *Artiodactyla*, which contains bovine and porcine species. The effect of phosphorylation at this site on BFSP2 function is unknown.

Phosphorylation of actin is known to play an essential role in the regulation of its depolymerization [56], though at the site Y53 rather than the ones noted in this study. Changes to the actin cytoskeleton are essential to fiber cell development [42], so it is likely that the changes in phosphorylation at the sites noted in the present study play an important role in regulating this process as well.

In total, the set of proteins showing changes in phosphorylation across the spatiotemporal gradient of the lens and the biological processes that they represent appear to tell the story that has been demonstrated by many other researchers: phosphorylation is a key regulator of lens fiber cell differentiation, resulting in significant changes to the behavior of critical lens proteins as lens fiber cells transition into their mature organelle-free state, though many open questions remain.

### 4.2. Oxidation Impacts Energy Metabolism and Major Lens Proteins

Given the known role of oxidation in the pathogenesis of age-related nuclear cataracts as a nonenzymatic and damaging modification that is associated with loss of protein solubility [57], the expectation in this study was that the presence of oxidized methionine would increase in the majority of cases when comparing the periphery of the lens to its more central regions. However, our results surprisingly show the opposite, with 16 out of the 18 discordantly altered oxidized peptides showing a decrease, rather than an increase, across the spatiotemporal gradient. There are several possible explanations for this. Given that these lenses were obtained from relatively young animals and that formation of methionine sulfoxide is a reversible modification, the pattern of oxidation here may be more indicative of where reactive oxygen species were primarily being generated in the lens at the time of tissue harvest rather than long term accumulation of oxidative damage. Lens epithelia and peripheral fiber cells contain mitochondria and are more exposed to oxygen in the aqueous humor, making them more susceptible to oxidation than the fiber cells in the organelle-free zone, which is heavily depleted in oxygen and where no aerobic respiration occurs [2]. Another possibility is that the oxidation noted in this study is artifactual and due to higher exposure of the outer layers of the lens to atmospheric oxygen compared to its inner layers during storage of the tissue. This is unlikely as lenses were stored within whole eyes at -80°C and only briefly following collection from the animals.

Regardless of the source, it is clear that oxidation preferentially affects certain types of proteins rather than uniformly impacting all proteins of the lens based upon the differing set of significant proteins and the biological pathways they map to in this analysis. Proteins relating to energy metabolism were much more represented in the oxidation data than the protein abundance or phosphorylation data. The reason for this is unclear. While it is expected that proteins involved in aerobic metabolism will tend to be more oxidized given their proximity to a major source of reactive oxygen species, the proteins in this case are related to glycolytic processes and should not be at greater risk for oxidation than any other cytosolic proteins.

Unlike most other proteins analyzed, α and β crystallins showed a discordant increase in oxidized peptides when comparing the periphery to the nucleus. This is in contrast to CRYZ, which does not share homology with α or β crystallins and functions as a quinone reductase [58]. This may be due to the α and β crystallins being among the most abundant and long-lived proteins in the lens, resulting in detectable accumulation of oxidative modifications across the spatiotemporal gradient even at this relatively young age.

### 4.3. Limitations and Future Directions

Because this study only examined lenses from pigs of approximately 6 months of age, we cannot determine how dissolution time differs based on age. It is likely that older lenses would require more time to dissolve and may not follow linear intervals as deeper layers have increased density. In future studies with lenses from other organisms, or in porcine lenses recovered from pigs over 6 months in age, new intervals will need to be established in order to evenly fractionate the lens. Additionally, although the methods presented here fractionated the lens into six sections, division into a larger number of sections is entirely feasible, allowing for examination of age-related changes in PTMs with even sharper resolution. Similarly, the age of the lenses used in this study are a limiting factor for the range of PTMs that can be assessed, given that many more age-related changes are likely to be detectable in tissue obtained from older animals. Future studies should focus on utilizing these techniques to analyze lenses from animals and human donors of advanced age to provide the greatest insight into the effects of aging on long-lived proteins in the lens.

## 5. Conclusions

The methods we have demonstrated in this study provide the ability to analyze age-related proteomic changes longitudinally using only endpoint lens tissue, providing an effective model for investigating changes across the spatiotemporal gradient. Here, we found several changes in oxidation and phosphorylation that modify essential lens proteins, such as crystallins and beaded filament structural proteins, as well as glycolytic enzymes which may play an important role in lens development and cataractogenesis. Moreover, our results demonstrate that there are detectable differences in abundance and PTMs of lens proteins from fractions of fiber cells that differ in age by only months.

## Author Contributions

Conceptualization, J.S.M., M.J.M., J.A.W.; methodology, J.S.M., O.K., M.R., G.E.M., M.J.M., J.A.W.; investigation, J.S.M., G.E.M., J.A.W.; formal analysis, O.K., M.R., G.E.M., M.J.M.; writing – original draft, J.S.M., J.A.W.; writing – review & editing, J.S.M., O.K., M.R., G.E.M., M.J.M., J.A.W.

## Funding

Funding for this research was provided by the University of Washington Nathan Shock Center of Excellence in the Basic Biology of Aging (NIA P30AG013280).

### Institutional Review Board Statement

Ethical review and approval were waived for this study due to all material being discarded tissue from livestock. No animals were raised for the purpose of this study. A permit to Obtain Specimens from Official Establishments from the North Carolina Department of Agriculture and Consumer Services Meat and Poultry Inspection Division was obtained prior to tissue collection.

### Data Availability Statement

All Skyline documents, raw files for quality control, and DIA data are available at Panorama Public. Access URL: https://panoramaweb.org/PigLen-sPTMs.url. ProteomeXchange ID: PXD040507.

## Appendix A

**Figure A1.**
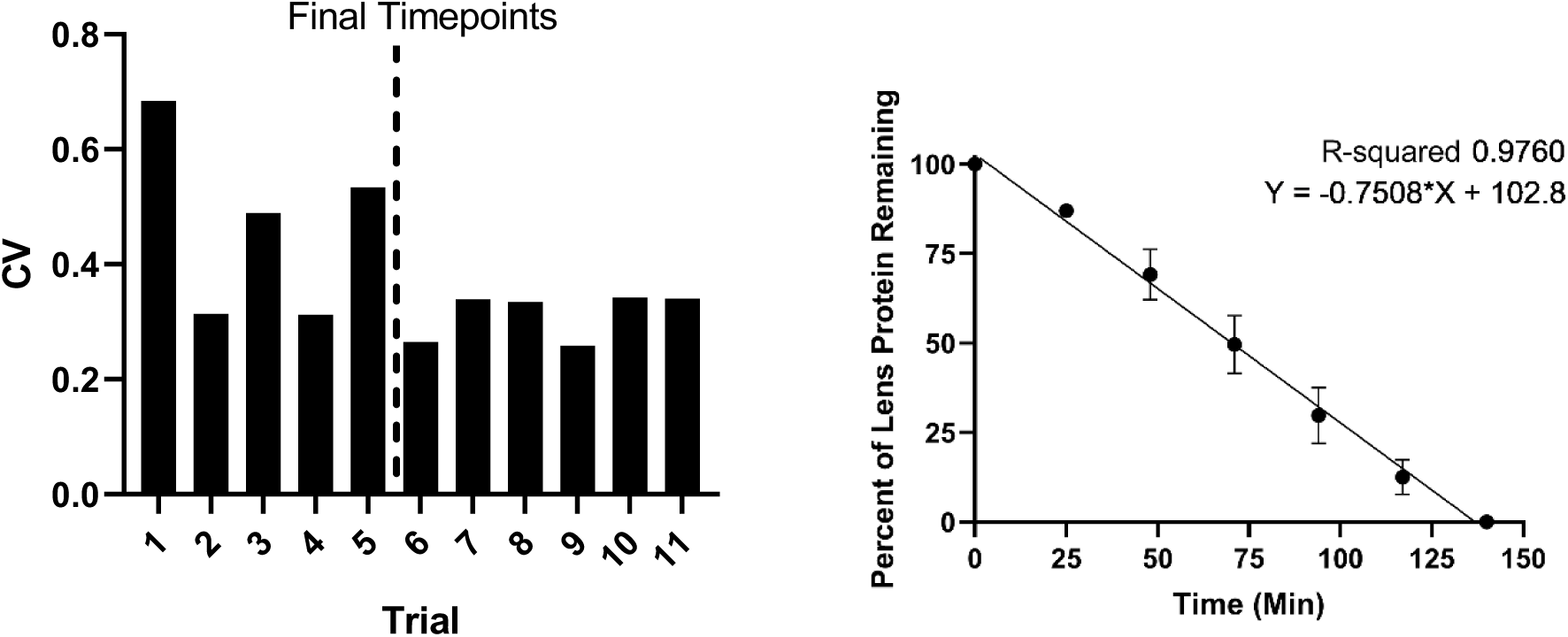
Dissolution Method Evenly Fractionates the Lens Across its Spatiotemporal Gradient. Porcine lenses were incubated in dissolution buffer to collect six successive layers of fiber cells. Left = Coefficients of variation (CV) of protein collected at across timepoints for 11 trials. Timepoints of 1, 2, 4, 8, 15, 30, 60, 90, and 120 minutes were used for the first trial, followed by assaying of protein content and use of linear regression to determine new time points that predicted even protein dissolution. This was repeated until CV remained relatively constant at ∼0.3 with the final timepoints of 25, 48, 71, 94, 117, and 140 minutes. Right = Linear regression of remaining lens protein at each timepoint using the final set of timepoints was assessed. N=6.

## Appendix B

Appendix B1: Full GEE and limma protein analysis dataset XLXS file

Appendix B2: Full GEE and limma peptide analysis dataset XLXS file

